# Comparative genomics guides elucidation of vitamin B12 biosynthesis in novel human associated *Akkermansia*

**DOI:** 10.1101/587527

**Authors:** Nina Kirmiz, Kadir Galindo, Karissa L. Cross, Estefani Luna, Nicholas Rhoades, Mircea Podar, Gilberto E. Flores

## Abstract

**Background:** *Akkermansia muciniphila* is a mucin-degrading bacterium found in the gut of most healthy humans and is considered a ‘next-generation probiotic.’ However, knowledge of the genomic and physiological diversity of human associated *Akkermansia* is limited, as only one species has been formally described.

**Results:** To begin to fill this knowledge gap, we reconstructed 35 high-quality metagenome assembled genomes from children and combined them with 40 other publicly available genomes from adults and mice for comparative genomic analysis. We identified at least four species-level phylogroups (AmI-AmIV) with distinct functional potentials. Most notably, we identified the presence of putative cobalamin (vitamin B12) biosynthesis genes within the AmII (n=26/28) and AmIII (n=2/2) phylogroups. To test these predictions, 10 novel strains of *Akkermansia* were isolated from adults and screened for essential vitamin B12 biosynthesis genes via PCR. Two strains of the AmII phylogroup were positive for the presence of vitamin B12 biosynthesis genes, while all AmI strains, including the type strain *A. muciniphila* Muc^T^, were negative. To demonstrate vitamin B12 biosynthesis, we measured the production of acetate, succinate, and propionate in the presence and absence of vitamin supplementation in representative strains of the AmI and AmII phylogroups since cobalamin is a cofactor in propionate metabolism. Results show that the *Akkermansia* AmII strain produced acetate and propionate in the absence of supplementation, which is indicative of *de novo* vitamin B12 biosynthesis. In contrast, acetate and succinate were the main fermentation products for the AmI strains when vitamin B12 was not supplied in the culture medium.

**Conclusions:** We identified *Akkermansia* strains as potentially important vitamin B12 biosynthetic bacteria in the human gut. This novel physiological trait of human associated *Akkermansia* may impact how these bacteria interact with the human host and other members of the human gut microbiome.

## BACKGROUND

*Akkermansia muciniphila* is a mucin degrading, gram-negative intestinal bacterium widely present in the human population, typically at 1 to 4% relative abundance [1, 2]. A number of studies in humans [3–5] and rodents [6–8] have found positive associations between its abundance and intestinal health, suggesting that *Akkermansia* may be a beneficial member of the gut microbiome and could be used as a biomarker of a healthy gut [9–11]. However, despite a diversity of phylotypes being reported by previous sequence-based studies, *A. muciniphila* Muc^T^ (ATCC BAA-835) represents the sole described species of the Verrucomicrobia phylum associated with humans [2, 12, 13]. Therefore, before we can fully assess the health potential of human associated *Akkermansia*, a comprehensive understanding of the genomic and physiological diversity of this lineage is needed.

Recently, a pangenomic study that included 33 new isolates from adults and 6 from laboratory mice provided insights into the population structure and evolutionary history of the *Akkermansia* lineage [14]. Specifically, this study revealed an open pangenome with at least three species-level phylogroups (AmI, AmII, and AmIII) that appear to be evolving independently. Although genomic differences amongst phylogroups were noted, the physiological consequences were not explored.

To continue to expand our understanding of the genomic content and functional potential of human associated *Akkermansia*, we reconstructed 35 *Akkermansia* genomes from children aged 2-9 years and combined our genomes with those from Guo et al. [14]. With these genomes, we identified novel diversity and several putative functional differences amongst the *Akkermansia* phylogroups. Most notably, we identified the presence of genes associated with *de novo* cobalamin (vitamin B12) biosynthesis in selected phylogroups of *Akkermansia*. Furthermore, using isolates obtained from healthy adults, we tested these genomic predictions and confirmed vitamin B12 biosynthesis by select strains of human-associated *Akkermansia*. These results build upon our understanding of the physiological capabilities of human-associated *Akkermansia* and demonstrate an important biosynthetic activity by this bacterial lineage that further expands its potential beneficial role in the intestinal environment.

## RESULTS

### Comparative genomics

A total of 334.9 Gbp of metagenomic sequence data was obtained from 70 children aged 2-9 years. Using SPAdes [15] to assemble contigs and MetaBAT [16] to bin contigs, we recovered 35 high quality metagenome assembled genomes (MAGs) of human associated *Akkermansia* (Table 1) from 35 of the 70 children. Completeness of the MAGs was relatively high, ranging from 68.5% to 95.5% with 31 of 35 MAGs > 90% complete. Likewise, contamination of the MAGs were low with all < 1%. On average, each MAG was 2.87 Mbp in length and contained approximately 2,420 protein-coding genes.

**Table 1.** Summary of 35 *Akkermansia* metagenome assembled genomes (MAGs) recovered from a diverse population of children aged 2-9 years living in Los Angeles, CA, USA. The genome of *A. muciniphila* Muc^T^ was not included in the averages presented at the bottom of the table

| Genome name | Phylogroup | Genome Properties |  |  |  | Assembly Properties |  |  |
| --- | --- | --- | --- | --- | --- | --- | --- | --- |
|  |  | Completeness (%) | Contamination (%) | Predicted Proteins | Coding Density (%) | Contigs | Length (Mbp) | GC Content (%) |
| *A. muciniphila* Muc<sup>T</sup> | AmI | 100 | 0.0 | 2,238 | 88.6 | 1 | 2.66 | 55.8 |
| CDI-75C-7 | AmI | 95.5 | 0.0 | 2,433 | 88.4 | 26 | 2.82 | 55.5 |
| CDI-92A-19 | AmI | 68.5 | 0.0 | 2,117 | 89.0 | 317 | 2.27 | 55.8 |
| CDI-93C-15 | AmI | 91.0 | 0.0 | 2,345 | 88.8 | 231 | 2.59 | 55.8 |
| CDI-16B-22 | AmI | 94.6 | 0.0 | 2,291 | 88.5 | 79 | 2.65 | 55.6 |
| CDI-158B-12 | AmI | 95.5 | 0.0 | 2,302 | 88.2 | 42 | 2.70 | 55.3 |
| CDI-50B-13 | AmI | 95.5 | 0.0 | 2,293 | 88.4 | 22 | 2.72 | 55.6 |
| CDI-51A-11 | AmI | 95.5 | 0.0 | 2,295 | 88.4 | 22 | 2.72 | 55.6 |
| CDI-28A-8 | AmI | 95.5 | 0.0 | 2,301 | 88.2 | 27 | 2.71 | 55.4 |
| CDI-85A-12 | AmI | 95.5 | 0.0 | 2,225 | 88.4 | 19 | 2.66 | 55.4 |
| CDI-30A-11 | AmI | 95.5 | 0.0 | 2,272 | 88.4 | 22 | 2.68 | 55.5 |
| CDI-42C-15 | AmI | 95.5 | 0.0 | 2,340 | 88.4 | 25 | 2.74 | 55.2 |
| CDI-151B-10 | AmI | 94.6 | 0.0 | 2,383 | 88.3 | 64 | 2.77 | 55.4 |
| CDI-193A-6 | AmI | 95.5 | 0.0 | 2,416 | 88.3 | 32 | 2.83 | 55.4 |
| CDI-143C-7 | AmII | 81.1 | 0.9 | 2,301 | 88.4 | 206 | 2.67 | 58.7 |
| CDI-10B-12 | AmII | 94.6 | 0.0 | 2,439 | 88.1 | 32 | 2.98 | 58.3 |
| CDI-128B-11 | AmII | 92.8 | 0.0 | 2,428 | 88.1 | 22 | 2.96 | 58.3 |
| CDI-129B-12 | AmII | 88.3 | 0.0 | 2,375 | 88.1 | 229 | 2.70 | 58.5 |
| CDI-77C-9 | AmII | 95.5 | 0.0 | 2,435 | 88.0 | 25 | 2.99 | 58.2 |
| CDI-24B-9 | AmII | 94.6 | 0.0 | 2,450 | 88.2 | 31 | 3.02 | 58.1 |
| CDI-182B-6 | AmII | 95.5 | 0.0 | 2,478 | 88.1 | 22 | 3.02 | 58.3 |
| CDI-198C-9 | AmII | 95.5 | 0.0 | 2,512 | 88.3 | 51 | 3.00 | 58.2 |
| CDI-69C-9 | AmII | 95.5 | 0.0 | 2,421 | 88.2 | 24 | 2.96 | 58.2 |
| CDI-138A-11 | AmII | 95.5 | 0.0 | 2,483 | 88.0 | 25 | 3.01 | 58.2 |
| CDI-70C-8 | AmII | 95.5 | 0.0 | 2,481 | 88.0 | 32 | 3.01 | 58.2 |
| CDI-26A-8 | AmII | 92.8 | 0.9 | 2,610 | 88.1 | 251 | 2.95 | 58.0 |
| CDI-34A-8 | AmII | 95.5 | 0.0 | 2,558 | 87.2 | 29 | 3.09 | 57.8 |
| CDI-65B-6 | AmII | 87.4 | 0.0 | 2,538 | 87.4 | 55 | 3.04 | 57.8 |
| CDI-203B-7 | AmII | 94.6 | 0.9 | 2,479 | 87.8 | 25 | 2.99 | 58.1 |
| CDI-150B-9 | AmIV | 95.5 | 0.0 | 2,457 | 87.7 | 29 | 2.99 | 57.2 |
| CDI-12C-16 | AmIV | 95.5 | 0.0 | 2,461 | 87.8 | 32 | 2.99 | 57.2 |
| CDI-156A-7 | AmIV | 95.5 | 0.9 | 2,532 | 87.5 | 56 | 3.05 | 56.7 |
| CDI-74B-7 | AmIV | 94.6 | 0.9 | 2,502 | 87.4 | 48 | 3.01 | 56.7 |
| CDI-18B-8 | AmIV | 94.6 | 0.0 | 2,509 | 88.0 | 124 | 2.95 | 56.9 |
| CDI-148A-8 | AmIV | 95.5 | 0.9 | 2,557 | 87.3 | 46 | 3.04 | 56.0 |
| CDI-13A-11 | AmIV | 95.5 | 0.0 | 2,670 | 88.1 | 66 | 3.20 | 56.6 |
| **AVERAGE** | **-** | **93.4** | **0.2** | **2,419.7** | **88.1** | **68.23** | **2.87** | **56.9%** |

To explore the genomic diversity of human associated *Akkermansia*, we performed a pangenomic analysis using tools in anvi’o [17, 18]. These analyses included the closed genome of the type strain [12] as well as 33 other human-associated and 6 mouse-associated *Akkermansia* genomes [14]. Previously, these 40 *Akkermansia* genomes were used to define three species-level phylogroups, AmI, AmII, and AmIII [14]. Merging our 35 MAGs with these 40 other genomes, we were able to regenerate the three phylogroups, but also observed a fourth phylogroup (AmIV) based on average nucleotide identity (ANI) calculated using PyANI [19] (heatmap in Figure 1). Phylogroup AmI, which includes the type strain *A. muciniphila* Muc^T^, contained the largest number of genomes with 40, followed by AmII (n=26), AmIV (n=7), and AmIII (n=2). Phylogroup AmIII was not observed in any of our 35 MAGs. Interestingly, both AmI and AmII included isolates obtained from mice. Within each phylogroup, ANI ranged from 93.94% to 99.98% across > 65% of each pair of genomes (Supplemental Figure 1). All between phylogroup ANI comparisons were < 92%. One genome in AmIV (CDI-148A-8) showed lower similarity (on average ∼94%) to other genomes within this phylogroup, possibly indicating further species level diversity within human associated *Akkermansia*. Across all phylogroups, we identified 6,557 gene clusters (GCs) with 1,021 found in all 75 genomes and 1,240 found in only one genome (Figure 1). Functional genes within the core included the cytochrome bd [20] (Amuc_1694 and Amuc_1695) and Type IV pili genes [21–23] (Amuc_1098 – Amuc_1102) previously characterized from *A. muciniphila* Muc^T^.

**Figure 1.**
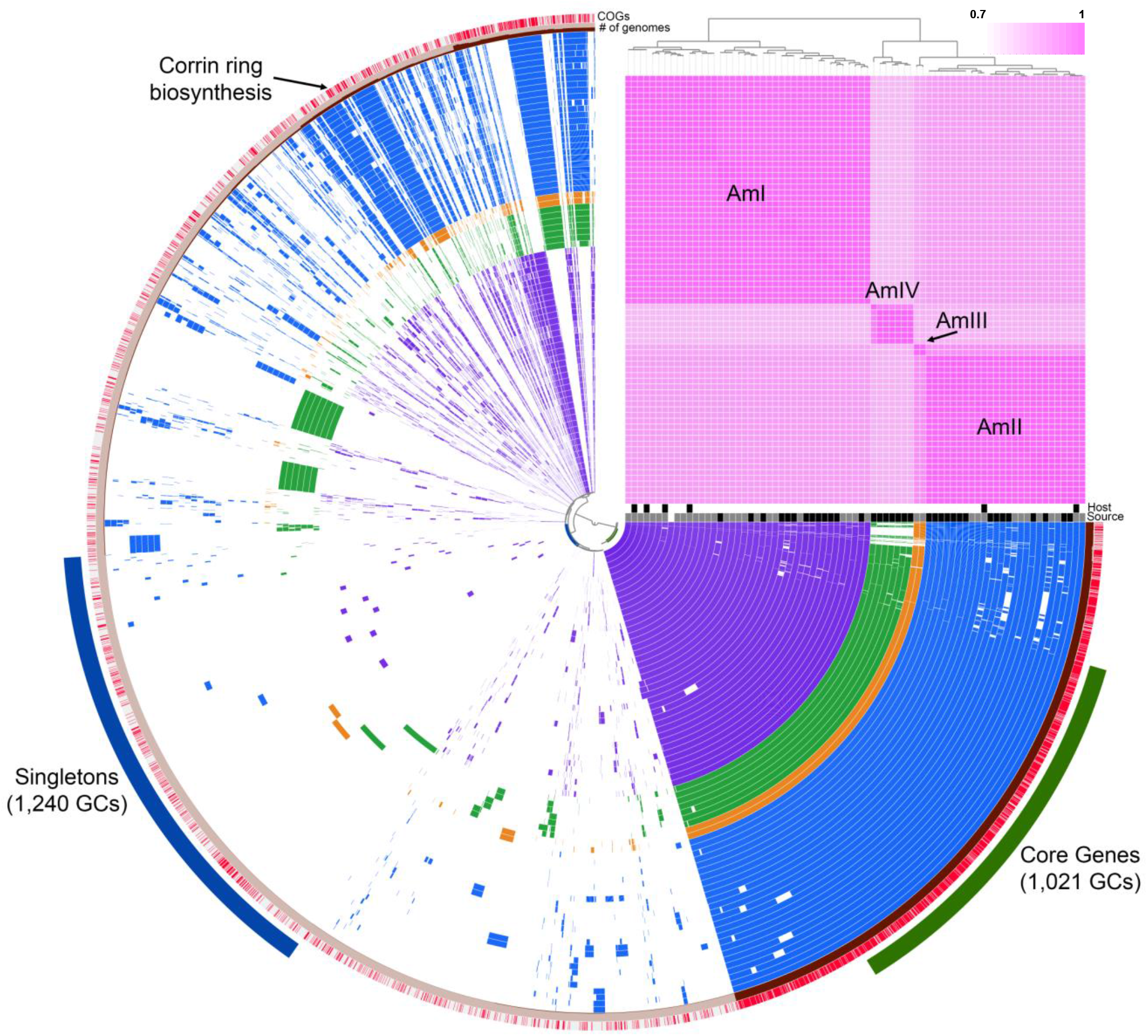
Pangenome of 75 Akkermansia genomes generated using anvi’o [17, 18]. Each concentric circle represents a bacterial genome with purple circles belonging to AmI, blue AmII, orange AmIII, and green AmIV phylogroups. Blank areas in each circle indicate absence of a particular gene cluster (GC) in that genome. A total of 6,557 GCs were observed across all genomes. Genomes are ordered by average nucleotide identity (ANI) as depicted by the pink heatmap in the upper right section of the figure. Host organisms are indicated just below the heatmap in white (human) or black (mouse) boxes. Similarly, genome sources are indicated in white [12], grey [14] and black (this work) boxes. The outer most ring is colored by presence (red) or absence (grey) of functional COG annotations. The next ring indicates the number of genomes that particular gene cluster was observed in. Singleton (blue) and core genes (green) are indicated outside of the concentric circles. Corrin ring biosynthesis genes are indicated in the AmII (blue) and AmIII (orange) genomes.

Next, we were interested in identifying functional gene predictions that differed amongst the phylogroups. Using Clusters of Orthologous Group (COG) annotations of GCs implemented in anvi’o, we observed 7 GCs putatively involved in the corrin ring stage of cobalamin (vitamin B12) biosynthesis within the AmII (n=24/26) and AmIII (n=2/2) phylogroups (Supplemental Data 1). To investigate these genes in greater detail, we manually inspected the annotations of all 75 genomes using Integrated Microbial Genome (IMG) [24] and Geneious 7.1.3 (https://www.geneious.com). With this approach, we confirmed the COG annotations and identified a cluster of 8 genes that appears to code for the corrin ring biosynthesis proteins in a subset of *Akkermansia* genomes (Figure 2). Included in this genomic region were genes *cbiK/X*, *cbiL*, *cbiC*, *cbiD*, *cbiET*, *cbiFGH*, and *cbiA* that encode the enzymes associated with the anaerobic pathway of corrin ring biosynthesis. This cluster also contains a gene annotated as a hypothetical protein, which shows some similarity to a putative cobalt transporter [25]. The content and arrangement of these genes was similar to the only other named species of the *Akkermansia* genus, *Akkermansia glycaniphila* Pyt^T^ previously isolated from a python [26]. Additionally, all 75 genomes contained most of the genes associated with the upstream (tetrapyrrole precursor biosynthesis: e.g. Amuc_0090 – 0091, Amuc_0417, Amuc_0896, and Amuc_1730) and downstream (nucleotide loop assembly, e.g. Amuc_1678 – Amuc_1683) stages of vitamin B12 biosynthesis [27]. Genes annotated as a TonB dependent transporter (e.g. Amuc_1684) and an extracellular solute-binding family 5 protein (e.g. Amuc_1685) that may be involved with vitamin B12 import were also identified adjacent to the nucleotide loop assembly genes in all but one genome.

**Figure 2.**
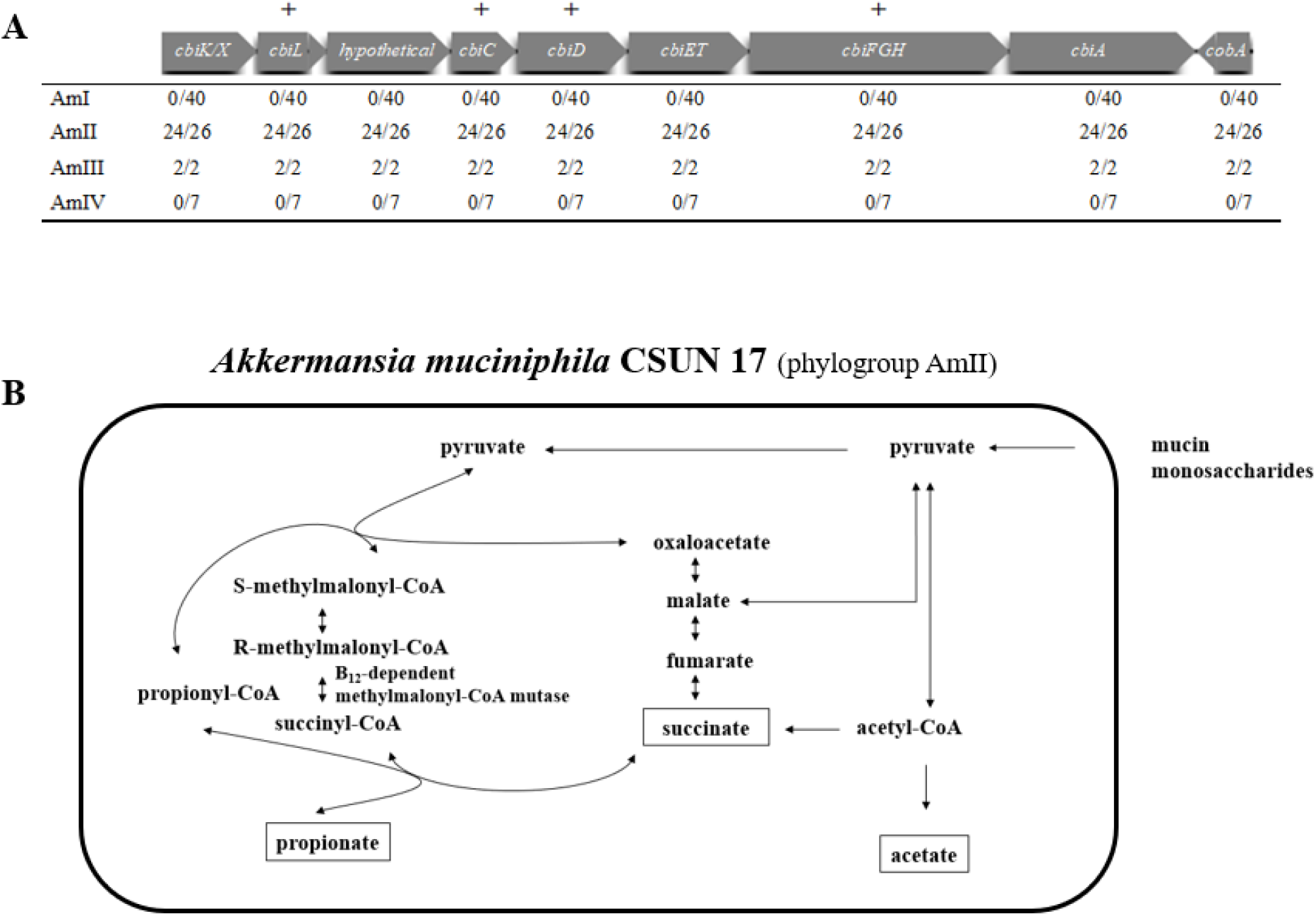
(A) Corrin ring biosynthesis gene cluster from isolate *A. muciniphila* CSUN-17 (phylogroup AmII). Presence of genes in the corrin ring biosynthesis gene cluster in phylogroups AmI, AmII, AmIII, and AmIV. Plus sign (+) above table indicates presence of gene in *A. muciniphila* CSUN phylogroup AmII isolates using PCR screen of *A. muciniphila* CSUN isolates. (B) Proposed strategy of propionate production in *A. muciniphila* CSUN-17 (phylogroup AmII) is shown involving *de novo* vitamin B12 biosynthesis leading to activation of methylmalonyl-CoA synthase and conversion of succinate to propionate.

*A. muciniphila* Muc^T^ was previously classified as a Cbi salvager because it lacked the genes coding for the enzymes to synthesize the corrin ring of vitamin B12, yet it needs this cofactor for methionine synthesis, nucleotide synthesis, queuosine synthesis and propionate metabolism [27]. Indeed, genes associated with these cellular functions were conserved across all phylogroups (Supplemental Data 1). Interestingly, the vitamin B12 independent methionine synthase II gene (*metE*) was present in 25 of 40 AmI genomes but not in any of the other genomes including the type strain Muc^T^. Together, these observations suggest that all *Akkermansia* examined here are able to acquire and likely remodel corrinoids from the environment for use, but some are also able to synthesize this important cofactor.

### Cultivation and validation of vitamin B12 biosynthesis

To determine if specific *Akkermansia* species/strains are indeed able to *de novo* synthesize vitamin B12, we first isolated several strains of *Akkermansia* from healthy adults and compared their near-full length 16S rRNA gene sequences with those from Guo et al. [14] in ARB [28] to determine phylogroup affiliation. Across phylogroups AmI, AmII, and AmIII, 16S rRNA gene sequences were all greater than 97% identical but nevertheless clustered into the known phylogroups (Supplemental Figure 2). Based on this approach, we identified eight AmI and two AmII isolates in our culture collection. Because our MAGs did not contain any full-length 16S rRNA gene sequences, we could not positively identify AmIV members among the isolates.

Next, using the AmII and AmIII genomes and the genome of *A. glycaniphila* Pyt^T^, we designed degenerate polymerase chain reaction (PCR) primers targeting four genes, *cbiL*, *cbiC*, *cbiD*, and *cbiFGH*, of the corrin ring biosynthesis gene cluster, which encode a cobalt-factor II C20-methyltransferase, a cobalt-precorrin-8 methylmutase, a cobalt-precorrin-5B (C(1))-methyltransferase, and a cobalt-precorrin-4 methyltransferase/precorrin-3B C17-methyltransferase, respectively (Supplemental Table 1). These genes were selected because they are predicted to give the best indication of cobamide production as described by Shelton et al. [27]. As expected, only isolates from the AmII phylogroup (CSUN-17 and CSUN-34) and *A. glycaniphila* Pyt^T^ showed positive amplification, whereas all AmI isolates (including *A. muciniphila* Muc^T^) failed to amplify (Table 2). Sequencing and BLASTing of these PCR amplicons from CSUN-17 against *A. glycaniphila* ERS 1290231 and *Desulfovibrio vulgaris* str. Hildenborough confirmed the identity of these gene fragments (Supplemental Table 2) clearly demonstrating the presence of select *cbi* genes in the AmII phylogroup.

**Table 2.**
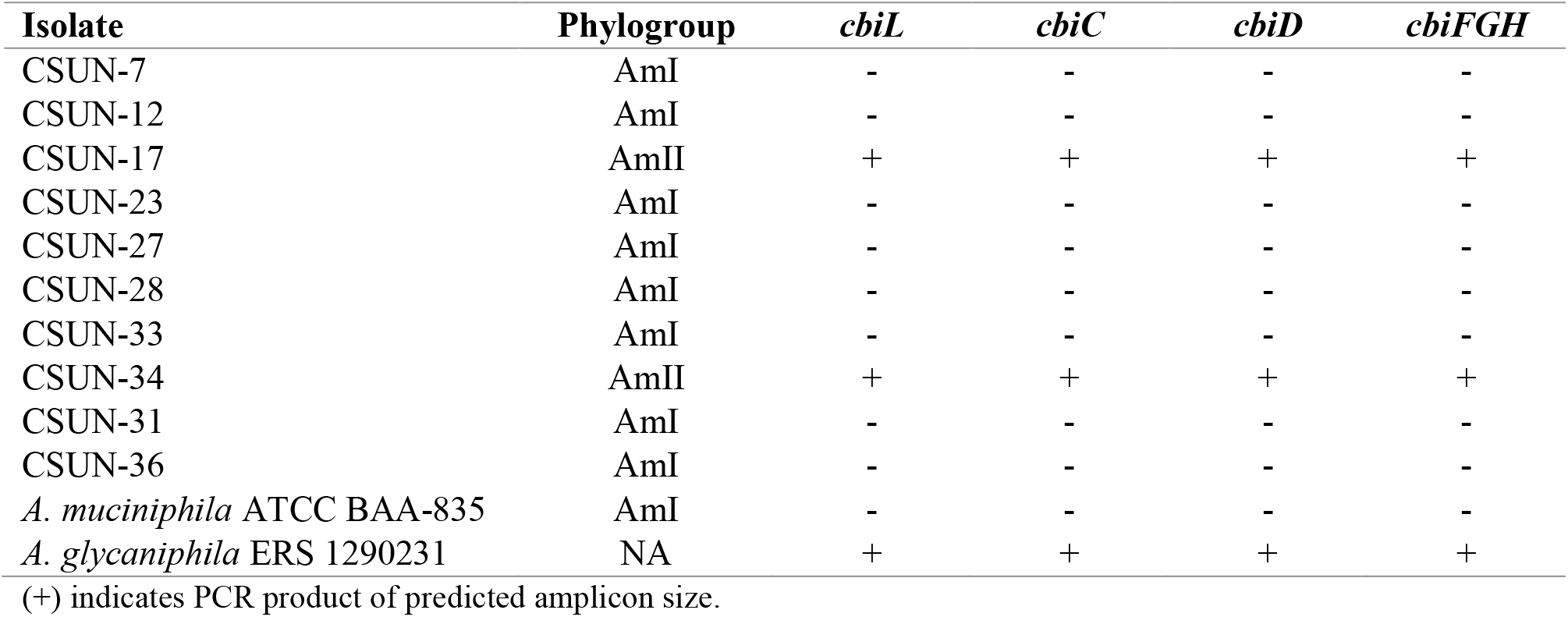
Presence of select corrin ring biosynthesis associated genes in CSUN *Akkermansia* isolates as determined by PCR.

It is known that many fermentative bacteria including *A. muciniphila* Muc^T^, use vitamin B12 to activate methylmalonyl-CoA synthase to convert succinate to propionate [29, 30]. Therefore, to demonstrate vitamin B12 biosynthesis *in vitro*, we quantified the production of succinate and propionate (and acetate) in the presence and absence of vitamin B12 in mucin medium (Figure 3). Our predictions were that the AmI phylogroup (represented by *A. muciniphila* Muc^T^) would produce acetate and succinate in the absence of vitamin B12, and acetate and propionate when B12 was present. For AmII, we predicted that acetate and propionate would be produced regardless of whether the culture medium was supplemented with vitamin B12. Results show that the AmI isolate produced propionate in a vitamin B12 concentration-dependent manner (Figure 3B,C). Also as expected, the CSUN-17 isolate (AmII), produced significant amounts of acetate and propionate in the absence and presence of vitamin B12, but production was more rapid with supplementation (Figure 3D-F). These results clearly indicate vitamin B12 biosynthesis by the *Akkermansia* species represented in the AmII phylogroup.

**Figure 3.**
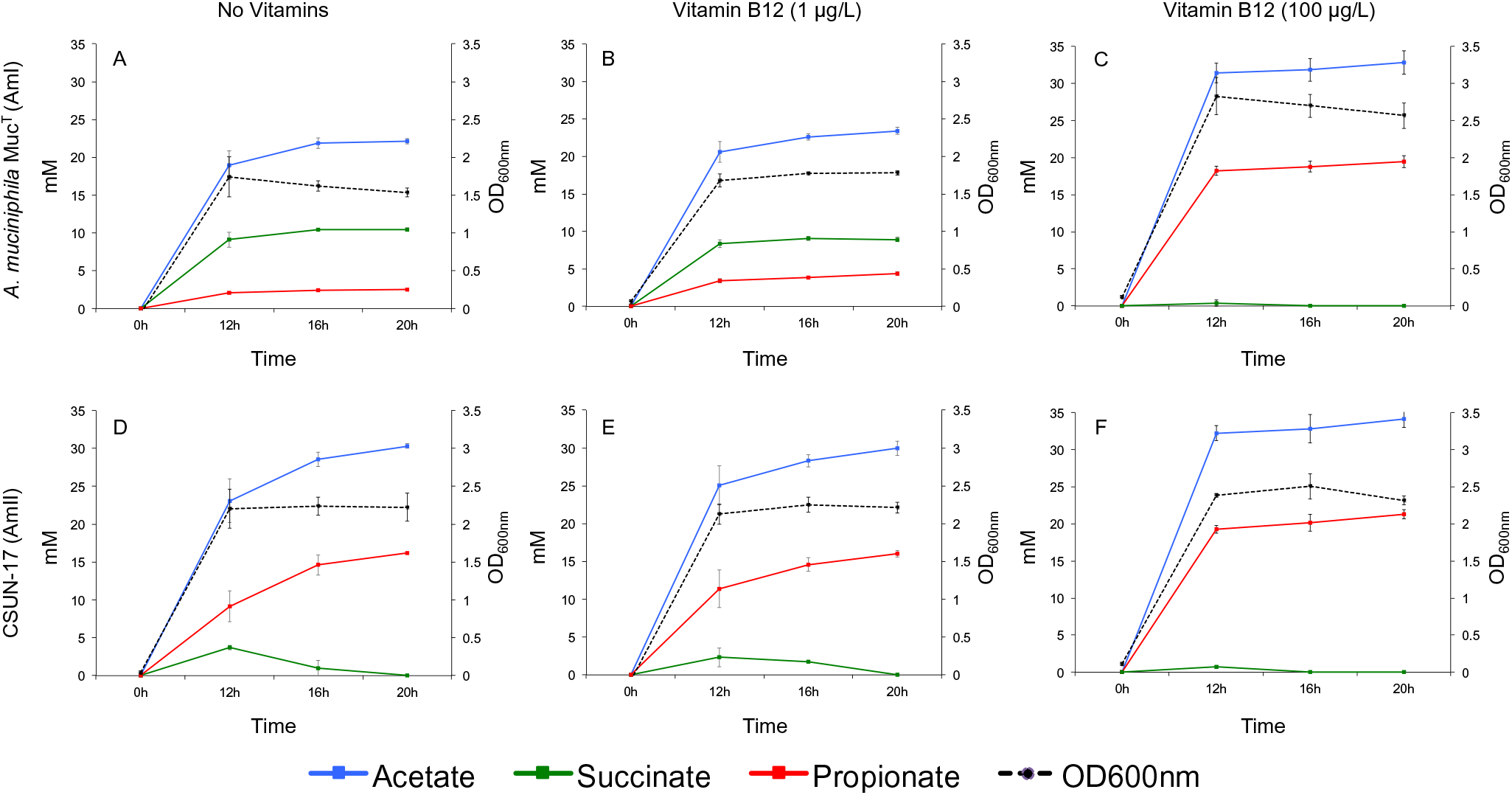
Production of acetate, succinate, and propionate through time by two strains of human associated *Akkermansia* grown on purified hog gastric mucin (1%) in the absence of vitamins (A, D), with ATCC MD-VS supplementation (B, E; 1% v/v vitamin solution/medium, vitamin B12 at a final concentration of 1 μg/L medium), and with vitamin B12 as cyanocobalamin (C, F; 100 μg/L). All values were averaged from four replicates and error bars represent the standard deviation. Background levels of organic acids present in the culture medium were subtracted from calculated averages when necessary.

## DISCUSSION

*A. muciniphila* is a common gut bacterium highly regarded as beneficial member of the human gut microbiome with important probiotic potential [10, 31]. Various studies have described positive associations between the abundance of *Akkermansia* and intestinal health [3, 4]. For example, *A. muciniphila* affects glucose metabolism, intestinal immunity, and its abundance in the gastrointestinal tract (GIT) is inversely correlated with diseases including Crohn’s disease, ulcerative colitis, and acute appendicitis [32–35]. Although a number of 16S rRNA gene variants have been observed [12] and dozens of isolates have been obtained [14], human-associated *Akkermansia* have largely been thought of as a single species and the functional potential beyond mucin degradation has gone largely unexplored. Here, we demonstrate that there are significant genomic and physiological differences amongst the human associated *Akkermansia*. Through comparative genomic analysis, we identified four phylogroups of human-associated *Akkermansia*, expanding the known genomic diversity of this lineage. Although all 16S rRNA gene sequenced examined here and elsewhere [32] are >97% identical, using an ANI of 95% across genomes as a species level delineation [36, 37] would suggest that each phylogroup represents a different species of *Akkermansia*. When we examined gene content, several phylogroup specific genes were identified that are predicted to code for functional differences amongst phylogroups further supporting species delineation. Most notably, we identified a complete set of genes involved in *de novo* biosynthesis of cobalamin, or vitamin B12, in two of the four phylogroups. We were able to validate these predictions *in vitro* using novel strains obtained from healthy adults. These findings demonstrate an ecological important function [38] not previously associated with human-associated *Akkermansia*, fundamentally altering our understanding of the diversity and physiology of this lineage. More broadly, these results continue to demonstrate the importance of merging next-generation sequencing approaches with traditional cultivation approaches to understand the basic biology of microorganisms of significance.

A recent comparative genomic analysis examining 11,000 bacterial genomes for cobamide production revealed that approximately 37% of bacteria are predicted to synthesize cobamides, yet 86% require them for at least one cellular function [27, 39]. Additionally, Degnan et al. found that most vitamin B12 dependent human gut bacteria lack the ability to synthesize vitamin B12 [39]. The type strain *A. muciniphila* Muc^T^ was included in the analysis by Shelton and colleagues [27] and was described as a Cbi salvager able to use exogenous sources of vitamin B12. Indeed, based on previous *in vitro* coculture experiments, *A. muciniphila* Muc^T^ can use at least three types of cobamides; cyanocobalamin supplied in culture medium, pseudovitamin B12 produced by *Eubacterium hallii* L2-7 [29], and an unknown form produced by *Anaerostipes caccae* [40]. Presumably, *Akkermansia* are able to import these various forms of cobalamin and use them directly or remodel the lower ligand to suit their needs. With our findings, some *Akkermansia* can now be considered producers of corrinoids, altering our understanding of how they interact with other members of the human gut microbiome and potentially their human host. However, questions remain regarding the type of cobalamin produced by AmII members and more generally regarding the specificity and efficiency of cobamide import and remodeling by all *Akkermansia*.

Cobalamin produced by bacteria and archaea in the large intestine is not readily available to the human host for two main reasons [38]. First, the receptors responsible for cobalamin absorption are found in the small intestine, which is not as densely colonized by bacteria as the large intestine in times of health. Second, although bacteria produce many different types of cobalamin, their contribution to the available pool of cobalamin is small because many of the forms produced by bacteria are not recognized by human receptors. Thus, bacteria are thought of more as competitors for dietary cobalamin than suppliers. However, if a bacterium colonized the small intestine and produced an appropriate form of cobalamin then the cofactor is possibly available to the human host. With regards to *Akkermansia*, we do not yet know the form of cobalamin produced by AmII members but *Akkermansia*-like organisms have been observed throughout the human gastrointestinal tract, including in the small intestine (reviewed in [32]). Interestingly, phylogenetic analyses consistently group AmII and AmIII isolates [14] with clones and other sequences previously observed in the small intestine [32]. Because our genomic sequence data and isolates were obtained from fecal samples, we could not determine if the different phylogroups colonize different segments of the gastrointestinal tract, but it is intriguing to speculate.

Although we do not yet know if humans can directly benefit from vitamin B12 produced by *Akkermansia*, there are indirect benefits resulting from the altered metabolites produced when vitamin B12 is available. Specifically, the type and quantity of short chain fatty acids (SCFA) produced during fermentation influences host health [41–43]. For example, propionate is known to help regulate appetite by stimulating the release of peptide YY (PYY) and glucagon like peptide-1 (GLP-1) by human colonic cells [44]. Less is known about the potential benefits of succinate in the human gut but in the mice cecum, succinate does improve glucose homeostasis via intestinal gluconeogenesis [45]. Conversely in the human small intestine, succinate has been shown to trigger a type 2 immune inflammatory response initiated by epithelial tuft cells [46]. Thus, possessing the ability to synthesize vitamin B12 *de novo* would suggest that the AmII and AmIII phylogroups have the potential to consistently produce more propionate than succinate during mucin fermentation and, as a result, influence gut epithelial cell behavior. If the AmII and/or AmIII phylogroups do colonize the small intestine, being able to consistently produce propionate over succinate could have significant health implications.

In addition to propionate metabolism, *Akkermansia* are predicted to use vitamin B12 as a cofactor for methionine biosynthesis using methionine synthase type I (MetH). All genomes possessed the *metH* gene; however, select AmI genomes (n=25/40) also contain the B12 independent methionine synthase II gene (*metE*), suggesting that these select AmI strains can generate methionine in the absence of vitamin B12. Given that the AmI phylogroup does not synthesize vitamin B12, this would allow production of this essential amino acid when exogenous corrinoids were unavailable. How readily available corrinoids are to *Akkermansia* either from other bacterial producers or host diet is unknown but possessing both variants may be an adaptive strategy for AmI strains.

*A. muciniphila* is being explored as a commercial probiotic and/or therapeutic agent [35]. Recent studies have reported large scale cultivation of *A. muciniphila* on a defined medium, safe for human consumption [22] and evaluated the stability and viability of the bacterium in dark chocolate [47]. Our results nevertheless indicate that there are still gaps in understanding the diversity and physiology of human associated Verrucomicrobia that need to be explored.

## CONCLUSION

Here we carried out a pangenomic analysis of 75 *Akkermansia* genomes and identified at least four species-level phylogroups (AmI-AmIV) with differing functional potentials. However, a polyphasic taxonomic characterization that includes robust phenotypic and genomic analyses is needed to verify species designations. Quantification of SCFA by select strains in the presence and absence of vitamin B12 supplementation demonstrated cobalamin biosynthesis by AmII strains. This work alters our understanding of how *Akkermansia* interacts with its human host and other members of the human gut microbiome in its unique environment. Future work will focus on other genomic similarities and differences identified in our analysis, but also continue to explore vitamin B12 production and acquisition using our culture collection. We are also continuing to isolate novel strains from healthy adults attempting to obtain representatives of each phylogroup observed or others that have yet to be observed.

## METHODS

### METAGENOMIC STUDIES

#### Recruitment and Sampling

Samples used for metagenomic sequencing were obtained from healthy children aged 2-9 years as described elsewhere (Herman et al., *in review*). These participants were consented under protocol #1314-223 approved by the Institutional Review Board (IRB) at California State University, Northridge (CSUN). Supplemental Data 2 provides unidentifiable demographic information of each child included in this study.

#### DNA extraction, Library prep, sequencing

Parents collected fecal samples in the privacy of their homes using sterile, double-tipped swabs by swabbing toilet paper (or diapers) after use. Samples were then frozen at −20 °C within 24 hrs of collection, and transported on blue ice to the lab (<30 min transit), where they were stored at −80 °C. This protocol is minimally invasive and has been successfully used in many similar, community-based research projects [48–50].

DNA was extracted from approximately ∼0.1g of collected samples using the MOBIO Power Soil^®^, DNA Isolation kit following a modified extraction protocol [51]. Extracts were then quantified using a Qubit 2.0 with high sensitivity reagents, and 100ng of DNA from each sample was sheared into 300bp fragments using a Covaris M220 [52]. The NEBNext® Ultra^™^ DNA Library Prep Kit for Illumina® [53] was used to prep dual indexed metagenomic libraries from the sheared samples. Libraries were confirmed using a BIO-RAD Experion^™^ Automated Electrophoresis System, and KAPA qPCR NGS library quantification. Two sequencing runs of the multiplexed libraries were conducted on an Illumina Hiseq 2000 (2 × 100bp) at the University of California Irvine, Genomics High-Throughput Facility.

#### Metagenomic Sequence processing

Raw fastq files from each sample were trimmed using TRIMMOMATIC [54] (ILLUMINACLIP:TruSeq3-PE.fa:2:30:10 LEADING:3 TRAILING:3 SLIDINGWINDOW:4:15 MINLEN:36). Trimmed sequences were then screened against the human genome (GRCh38) using DeconSeq [55] in order to remove any potential human DNA sequences. Non-human sequences were further cleaned using PRINSEQ [56] with the following parameters: - min_qual_mean 20 and -ns_max_n 3. Remaining sequences without a matepair were removed and paired sequences were assembled using the default parameters for metagenomes in SPAdes [15]. Resulting contigs > 2 Kbp were binned used MetaBAT [16] with default parameters and the taxonomy and completeness of bins were verified using the taxonomy workflow of CheckM [57] against the Verrucomicrobia phylum. We determined that bins confidently identified as ‘k_Bacteria (UID2982)’ were *Akkermansia* and evaluated the quality of those bins further using MiGA [58]. Assembled contigs (>2kb) from each child with a high quality *Akkermansia* bin were submitted to IMG-M where they were annotated using their workflow [24]. Both IMG and Geneious 7.1.3 (https://www.geneious.com) were used to manually inspect annotations of interest. Vitamin B12 associated genes were detected by searching for annotations from Enzyme Commission (EC) numbers, IMG terms, pfam, and Clusters of Orthologous Groups (COG) [24, 59, 60]. The annotations included those used by Shelton et al. [27] and Degnan et al. [39].

#### Pangenome analysis

To explore the *Akkermansia* pangenome, we combined our 35 MAGs with 40 other publically available genomes in anvi’o [17, 18]. Assembled fasta files were first converted to ‘db’ files using the ‘anvi-script-FASTA-to-contigs-db’ command that uses Prodigal [61] to call open reading frames. Each ‘db’ file was then annotated against the COG database [62] using ‘anvi-run-ncbi-cogs’ with the ‘--use-ncbi-blast’ flag. After generating the genome storage file with ‘anvi-gen-genomes-storage,’ the ‘anvi-pan-genome’ command was run with the identical parameters (--num-threads 12, --minbit 0.5, --mcl-inflation 10, --use-ncbi-blast) outlined in Delmont and Eren [17, 63, 64]. The pangenome was visualized and aesthetics were modified using the ‘anvi-display-pan’ command. To calculate average nucleotide identity (ANI) in anvi’o, the ‘anvi-compute-ani’ command, which utilizes PyANI [19], was used. To identify functions (i.e. COG annotations) that were differentially distributed amongst the phylogroups, we used the ‘anvi-get-enriched-functions-per-pan-group’ with phylogroups (AmI-AmIV) as the category.

### CULTIVATION STUDIES

#### Recruitment and Sampling

Fecal samples used in culturing of *Akkermansia* isolates were obtained from healthy adults using swabs as previously described [49] under IRB protocol #1516-146. Collected samples were refrigerated (4 °C) and transferred to culture medium (see below) within 24-hours of collection.

#### Enrichment, isolation, genomic DNA extraction, and 16S rRNA gene sequencing

Anaerobic mucin medium was modified slightly from Derrien et al. [13] and contained (l^-1^) 0.4 g KH_2_PO_4_, 0.53 g Na_2_HPO_4_, 0.3 g NH_4_Cl, 0.3 g NaCl, 0.1 g MgCl_2_ · 6H_2_O, 0.4 g NaHCO_3_, 1 mg resazurin, and 10 ml trace mineral solution as described by Ferguson and Mah [65]. pH of the medium was adjusted to 6.5. Medium was prepared with boiled MiliQ water under constant gassing with a gas mixture consisting of N_2_/CO_2_ (80:20, v/v). Culture medium was later modified to include 1mM L-threonine and 10 g/L tryptone (Oxoid) as described previously [30]. Broth medium was prepared in serum tubes or bottles and sealed with butyl rubber stoppers and aluminum crimp caps prior to autoclaving at 121°C and 15 psi for 15 minutes. Prior to inoculation, medium was reduced with autoclaved 0.05% Na_2_S · 9H_2_O and supplemented with 0.5% - 1.0% purified hog gastric mucin (Type III, Sigma-Aldrich, St. Louis, MO). Purified mucin was prepared by first autoclaving a 5% or 10% solution prepared in 0.01 M phosphate buffer (stock = 88.46 g/L KH_2_PO_4_ and 60.97 g/L K_2_HPO_4_), performing dialysis using a 12-14 kD membrane (Spectra/Por 4, Spectrum Laboratories, Rancho Dominguez, CA), centrifuging twice for 10 min at 10,000 rpm, and filter sterilizing through 0.2 μm syringe filters (Whatman GE Healthcare Life Sciences, Chicago, IL) into growth medium. For solid medium, Noble Agar (Difco, Detroit, MI) was added and plates were poured in an anaerobic chamber (Bactron IV, Sheldon Manufacturing, Inc., Cornelius, OR) under an atmosphere of N_2_/CO_2_/H_2_ (80/15/5, v/v). All incubations were performed at 37 °C in the Bactron IV anaerobic chamber.

Enrichments cultures targeting mucin-degrading bacteria were initiated by transferring fecal swabs into 5ml of anaerobic mucin medium in serum tubes and performing ten-fold serial dilutions up the 10^−7^. Cultures were incubated for up to 5 days, monitored daily for changes in turbidity, and inspected using phase-contrast microscopy (Zeiss Axioskop). Positive cultures with oval cells in pairs were further diluted in broth medium and/or transferred to solid medium until purity could be verified microscopically and by sequencing of the 16S rRNA gene. For sequencing, genomic DNA was isolated using the MoBio Ultraclean Microbial DNA Isolation Kit (Mo Bio, Carlsbad, CA) following the manufacturer’s instructions. Briefly, 1.8 mL of overnight bacterial culture was centrifuged at 10,000 × *g* for 30 seconds, the pellet was resuspended in 300 μL of MicroBead Solution (Mo Bio, Carlsbad, CA), and the DNA was subsequently isolated following the manufacturer’s instructions. For amplification of the 16S rRNA gene via PCR, 2 μL of extracted genomic DNA was added to 25 μL of GoTaq Green Master Mix (Promega, Madison, WI) and 1 μL of 10 μM universal primers 8F (5’- AGAGTTTGATCCTGGCTCAG-3’) and 1492R (5’-TACGGTTACCTTGTTACGA-3’) using a 50 μL final PCR reaction volume. PCR was conducted using an Eppendorf Mastercycler Pro S 96 well thermocycler using a program of an initial denaturation at 95°C for 3 min, followed by 30 cycles of 95°C for 45 sec, annealing at 45°C for 1 min, 72°C for 1 min, a final extension of 72°C for 7 min, and holding at 4°C. PCR reactions were purified using the QIAquick PCR Purification Kit (Qiagen). Initial sequencing of the 16S rRNA gene was performed using either the 8F or 1492R primer on an ABI Prism 3730 DNA sequencer (Laragen Sequencing and Genotyping in Culver City, CA). If cultures were pure and positively BLASTed to *A. muciniphila*, the near full length 16S rRNA gene was sequenced with additional primers (515F (GTGCCAGCMGCCGCGGTAA), 806R (GGACTACHVGGGTWTCTAAT), and 8F or 1492R). Sequences associated with each isolate were then assembled in Geneious 7.1.3 (https://www.geneious.com) and imported into ARB [28] as discussed below. General demographic information about donors is provided as Supplemental Table 3.

#### 16S rRNA gene phylogeny

To determine phylogroup affiliation of our isolates, 16S rRNA gene sequences of the Guo et al. [14] isolates were first extracted from their genomic sequence data and imported into ARB [28]. Once in ARB, gene sequences were aligned with secondary structure constraints against the 16S rRNA gene sequence of *A. muciniphila* Muc^T^, manually inspected, and those that were < 1000 bp were discarded. Similarly, 16S rRNA gene sequences of our novel isolates were imported and aligned in ARB. A custom alignment mask excluding nucleotide positions found in less than half of all isolates was generated and masked alignments were imported into MEGA7 [66] where phylogenetic reconstruction was generated using the maximum-likelihood approach. Because we knew the affiliation of the Guo et al [14] isolates, we were able to place our isolates in this framework based on placement in the 16S rRNA gene tree.

#### Corrin biosynthesis PCR screen of isolates and gene sequencing

To amplify conserved regions of corrin biosynthesis associated genes, degenerate primers were designed (Supplemental Table 1). Select corrin biosynthesis associated homologous sequences were aligned using BioEdit Sequence Alignment Editor Version 7.0.5 (http://www.mbio.ncsu.edu/BioEdit/page2.html) (locus tags of sequences used in the alignments are shown in Supplemental Table 4). All gene sequences were obtained from JGI IMG/ER. Conserved regions were found using the Accessory Application ClustalW Multiple alignment tool in BioEdit [67]. For amplification of corrin biosynthesis genes *cbiL*, *cbiC*, *cbiD*, and *cbiFGH*, 1uL of genomic DNA was added to 12.5 μL of GoTaq Green Master Mix (Promega, Madison, WI) and 1 μL of each 10 μM primer using a 25 μL final PCR reaction volume. PCR conditions were optimized, and a PCR screen of isolates was carried out in duplicates using a PCR program of an initial denaturation at 95°C for 2 min, followed by 25-35 cycles of 95°C for 45 sec, annealing at 52°C to 62°C for 30 sec to 1 min, 72°C for 45 sec, a final extension of 72°C for 5 min, and holding at 4°C. PCR amplicons were separated and visualized using 1% agarose gel. PCR products were purified using the QIAquick PCR Purification Kit (Qiagen). For the amplification of *cbiFGH*, the amplicon was excised out of the gel and gel purified using the PureLink Quick Gel Extraction and PCR Purification Combo Kit (Invitrogen). The amplicons were sequenced as described above. BioEdit Version 7.0.5 was used to analyze the sequences. Sequences of PCR amplicons from CSUN-17 were checked by blastx using the IMG and the NCBI database to examine similarity to vitamin B12 associated genes from the genomes *A. glycaniphila* ERS 1290231 and *Desulfovibrio vulgaris* str. Hildenborough (Supplemental Table 2).

#### Quantification of short chain fatty acids via HPLC

To quantify production of short-chain fatty acids with and without vitamin supplementation *A. muciniphila* Muc^T^ (AmI) and CSUN-17 (AmII) were grown in anaerobic mucin medium supplemented with 1mM L-threonine, 10 g/L tryptone (Oxoid), 1% purified mucin and vitamin supplementation depending on treatment conditions. For vitamin supplementation, we first performed the experiment using the ATCC MD-VS at the recommended concentration (10 ml/L). Because the concentration of vitamin B12 in the formulation is 100 fold less than those reported by Belzer et al. [29], we subsequently performed a second experiment with pure vitamin B12 (Sigma-Aldrich, St. Louis, MO) using a final concentration of 100 ng/ml. For all experiments, overnight cultures were transferred in appropriate medium 3 times with the final transfer used to inoculate 25ml of medium at 5% in quadruplicate for each isolate and treatment. OD_600nm_ (Eppendorf BioPhotometer plus) was recorded at inoculation and at 12, 16, and 20 hours. An additional 1.25 ml of culture was removed at each time point, centrifuged at 15,000 × g for 10 minutes, and the cell-free supernatant was filtered through a 13mm, 0.2μm SPARTAN HPLC syringe filter. Samples were stored at −20°C until HPLC analysis.

High-performance liquid chromatography (HPLC) was performed using a Waters Breeze 2 system (Waters Corp., Milford, MA, USA) equipped with a refractive index detector (model 2414). An Aminex HPX-87H column (Bio-Rad Laboratories) was used to measure production of short chain fatty acids (SCFA). Sulfuric acid (5mM) was used as the mobile phase at a flow rate of 0.6 mL/min. Peak areas and retention times were compared against known standards. Samples were also compared against a media-only control to determine background levels of acetate, propionate, and succinate present in the starting medium before growth. Approximately 3mM propionate was detected in the culture medium and subtracted from all respective measurements.

## LIST OF ABBREVIATIONS

GIT: Gastrointestinal tract
SCFA: short chain fatty acids
PCR: polymerase chain reaction
HPLC: high-performance liquid chromatography
MAGs: metagenome assembled genomes
ANI: average nucleotide identity
GC: gene cluster
COG: Clusters of Orthologous Groups
EC: Enzyme Commission
IMG: Integrated Microbial Genomes

## DECLARATIONS

### Ethics approval and consent to participate

Aspects of this work that included human subjects was approved by the Institutional Review Board at California State University, Northridge under protocol #1314-223 (metagenomic study) and #1516-146 (cultivation study). For the metagenomic study, verbal assent was obtained from each child and written consent was obtained from one parent/guardian. Written consent was obtained from each subject in the cultivation studies.

### Consent for publication

Not applicable.

### Availability of data and material

Genomic sequence data from Guo et al. [14] is available at GenBank under BioProject #PRJNA331216. Our quality filtered metagenomic sequence data are available at GenBank under BioProject #PRJNA525290. Additionally, assembled conitgs >2kb from children who contained an *Akkermansia* bin are available in IMG under GOLD Study ID Gs0133482. It is important to note that contigs available in IMG include not only *Akkermansia* contigs, but also all contigs from each child. Supplemental Data 3 has a list of the *Akkermansia* contigs in IMG that were included in our analysis. Near full-length 16S rRNA gene sequences of our isolates are available in GenBank under accession numbers MK577303 – MK577312. Corrin gene sequences of isolate CSUN-17 are available in GenBank under accession numbers MK585566 – MK585569.

### Authors’ contributions

NK conducted wet lab work, analyzed and interpreted the data, and wrote the paper. KG performed bioinformatic analysis, and analyzed and interpreted the data. KLC conducted wet lab work and analyzed and interpreted the data. EL collected samples, conducted wet lab work, and analyzed and interpreted the data. NR conducted wet lab work and analyzed and interpreted the data. MP analyzed and interpreted the data. GEF conceived of and designed the study, performed bioinformatics analysis, analyzed and interpreted the data, and wrote the paper. All authors read and approved the final manuscript.

## Acknowledgements

The authors would like to thank Dara Fluke, Erik Hearn, Christopher Herrera, Arinnae Kurdian, Jonathan Lemus, Claudia Mendoza, and Priscilla Salcedo for help in the isolations. We would also like to thank Dena Herman, Joan Maltese, Zelzah Guzman, Alison Gambou, Ann-Marie Pham, Griseida Ruiz, and Jessica Saavedra for all their efforts in participant recruitment for the metagenomic studies. Finally, thanks to all study participants that provided fecal samples, without you none of this would be possible.

---

**Supplemental Table 1.**
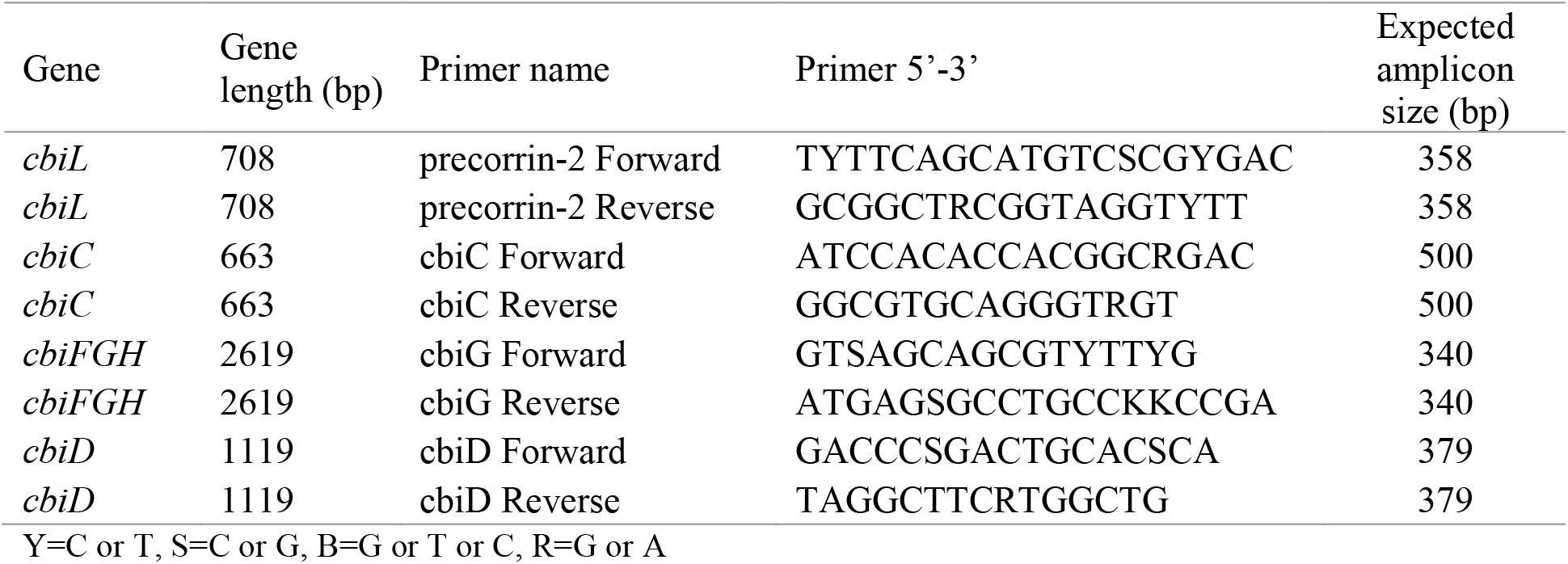
Corrin ring biosynthesis degenerate primers.

**Supplemental Table 2.** Corrin ring biosynthesis sequenced amplicons from CSUN-17 homology to *A. glycaniphila* and *Desulfovibrio vulgaris*.

| Gene name | *A. glycaniphila* gene name<sup>a</sup> | Percent identity to *A. glycaniphila*<sup>b</sup> | *Desulfovibrio vulgaris* gene name<sup>a</sup> | Percent identity to *Desulfovibrio vulgaris*<sup>b</sup> |
| --- | --- | --- | --- | --- |
| *cbiL* | precorrin-2 C20-methyltransferase /cobalt-factor II C20-methyltransferase | 65% | precorrin-2 C20-methyltransferase | 48% |
| *cbiC* | precorrin-8X methylmutase | 80% | precorrin-8X methylmutase | 49% |
| *cbiD* | cobalt-precorrin-5B (C1)-methyltransferase | 70% | cobalamin biosynthesis protein CbiD | 46% |
| *cbiFGH* | cobalt-precorrin 5A hydrolase/precorrin-3B C17-methyltransferase | 56% | precorrin-4 C11-methyltransferase | 32% |

<sup>a</sup>IMG/ER gene names using BLASTx against *A. glycaniphila* ERS 1290231 and *Desulfovibrio vulgaris*
(Hildenborough)
<sup>b</sup>Percent identity of CSUN-17 sequence using BLASTx against *A. glycaniphila* ERS 1290231 and *Desulfovibrio vulgaris* (Hildenborough)

**Supplemental Table 3.**
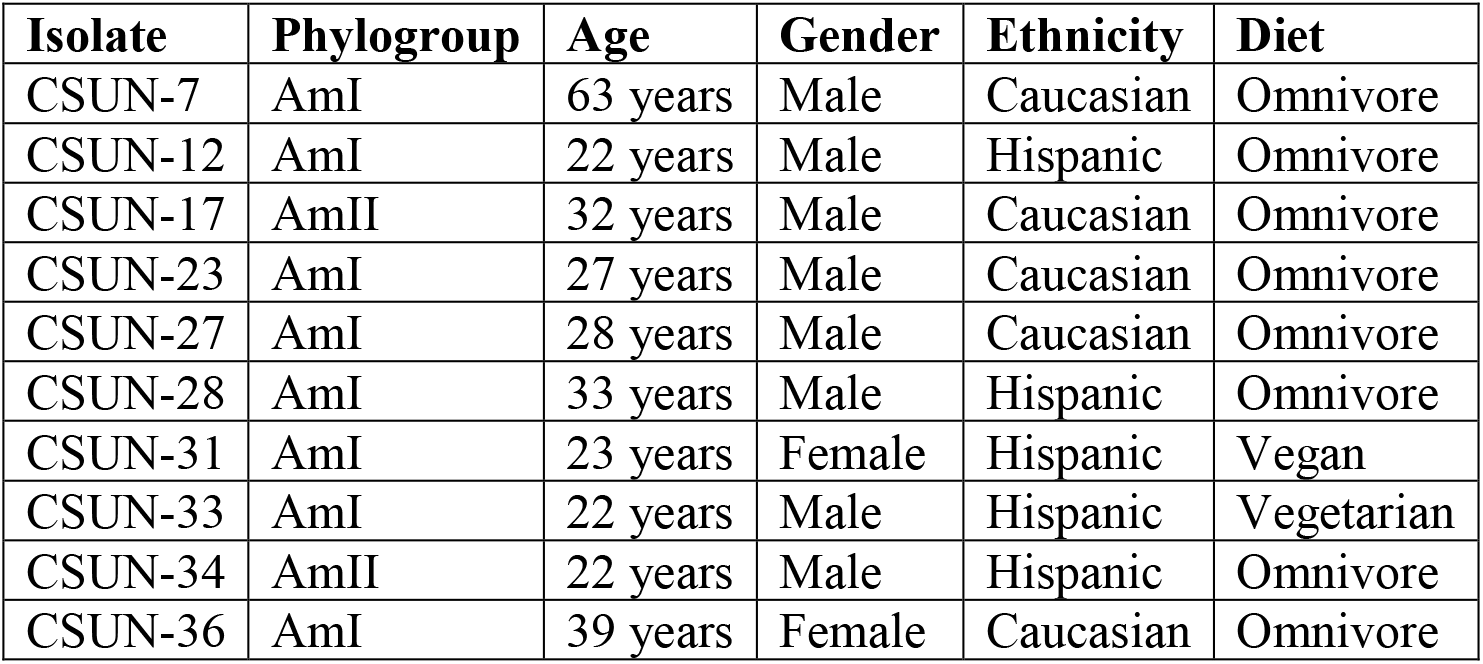
Demographic information of individuals who provided fecal samples from which strains of *Akkermansia* were isolated.

| Isolate | Phylogroup | Age | Gender | Ethnicity | Diet |
| --- | --- | --- | --- | --- | --- |
| CSUN-7 | AmI | 63 years | Male | Caucasian | Omnivore |
| CSUN-12 | AmI | 22 years | Male | Hispanic | Omnivore |
| CSUN-17 | AmII | 32 years | Male | Caucasian | Omnivore |
| CSUN-23 | AmI | 27 years | Male | Caucasian | Omnivore |
| CSUN-27 | AmI | 28 years | Male | Caucasian | Omnivore |
| CSUN-28 | AmI | 33 years | Male | Hispanic | Omnivore |
| CSUN-31 | AmI | 23 years | Female | Hispanic | Vegan |
| CSUN-33 | AmI | 22 years | Male | Hispanic | Vegetarian |
| CSUN-34 | AmII | 22 years | Male | Hispanic | Omnivore |
| CSUN-36 | AmI | 39 years | Female | Caucasian | Omnivore |

**Supplemental Table 4:** Locus tags of sequences used for corrin ring biosynthesis degenerate primer design.

| Gene | Locus tag |
| --- | --- |
| *cbiL* | Ga0175004_112520<br>T370DRAFT_00397<br>Ga0256628_10023618<br>Ga0257029_1018451<br>Ga0257030_10082613<br>Ga0257062_100012256<br>Ga0257032_118796<br>Ga0257040_100005132<br>Ga0257041_10019915<br>Ga0257042_100005192<br>Ga0257043_10000156<br>Ga0257044_106873<br>Ga0257047_1015355<br>Ga0257051_1033411<br>Ga0257052_10000756<br>Ga0257053_10015516<br>Ga0257060_10006251 |
| *cbiC* | Ga0175004_112518<br>DESPIG_00351<br>Ga0256628_10023616<br>Ga0257029_1018453<br>Ga0257030_10082611<br>Ga0257062_100012254<br>Ga0257032_118794<br>Ga0257040_100005134<br>Ga0257041_10019913<br>Ga0257042_100005190<br>Ga0257043_10000158<br>Ga0257044_106871<br>Ga0257047_1015357<br>Ga0257051_1033413<br>Ga0257052_10000758<br>Ga0257053_10015514<br>Ga0257060_10006249 |
| *cbiG* | Ga0175004_112515<br>HMPREF0179_00024<br>Ga0256628_10023613<br>Ga0257029_1018456<br>Ga0257030_1008268 |
|  | Ga0257062_100012251<br>Ga0257032_118791<br>Ga0257040_100005137<br>Ga0257041_10019910<br>Ga0257042_100005187<br>Ga0257043_10000161<br>Ga0257044_108973<br>Ga0257047_1015360<br>Ga0257051_1033416<br>Ga0257052_10000761<br>Ga0257053_10015511<br>Ga0257060_10006246 |
| *cbiD* | Ga0175004_112517<br>HMPREF0178_02483<br>Ga0256628_10023615<br>Ga0257029_1018454<br>Ga0257030_10082610<br>Ga0257062_100012253<br>Ga0257032_118793<br>Ga0257040_100005135<br>Ga0257041_10019912<br>Ga0257042_100005189<br>Ga0257043_10000159<br>Ga0257044_108971<br>Ga0257047_1015358<br>Ga0257051_1033414<br>Ga0257052_10000759<br>Ga0257053_10015513<br>Ga0257060_10006248 |

