## Supplementary figures and images for "Comparative genomics guides elucidation of vitamin B12 biosynthesis in novel human associated *Akkermansia*"

### Supplemental figures

#### Slide 1
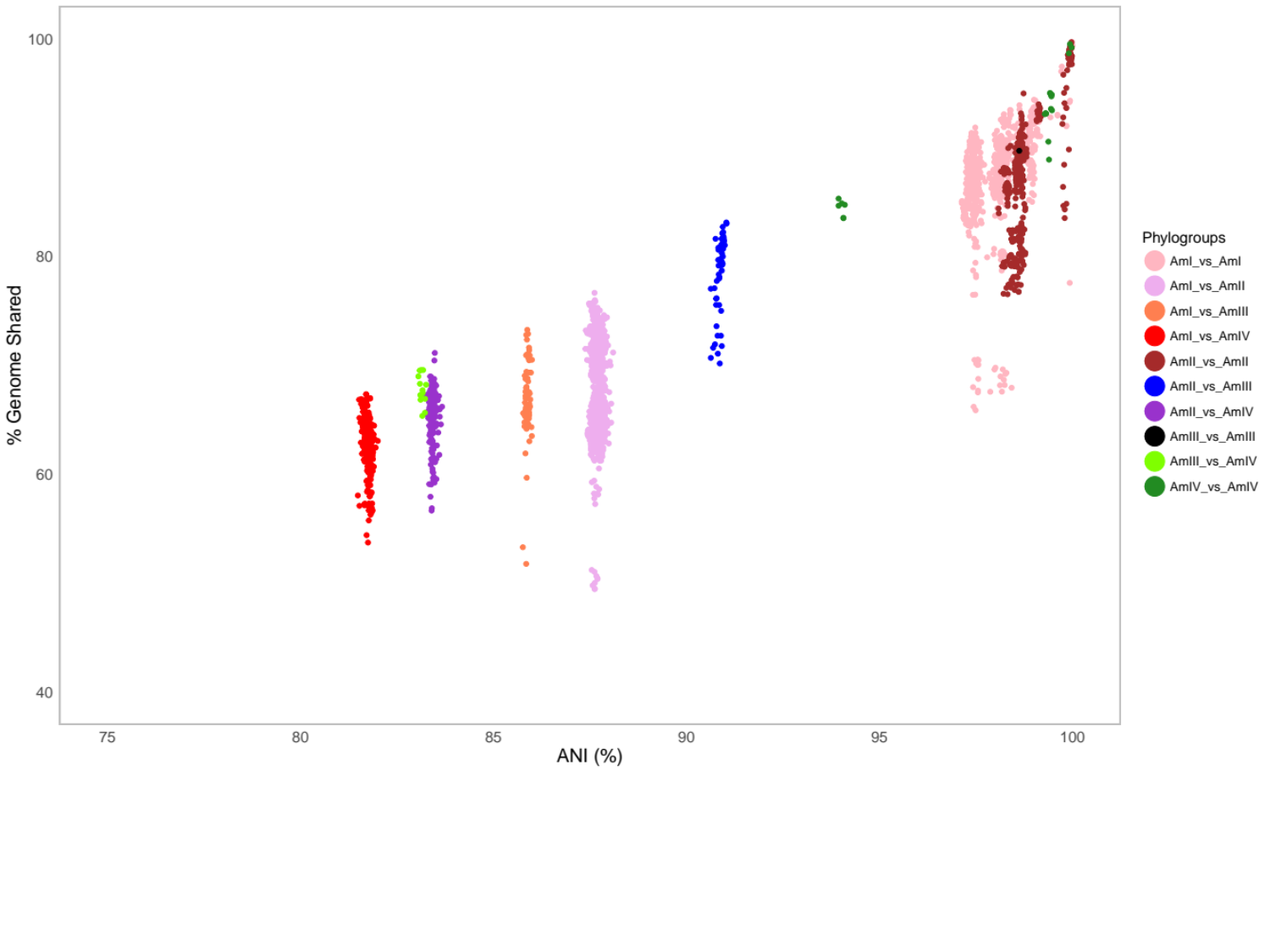

#### Slide 2
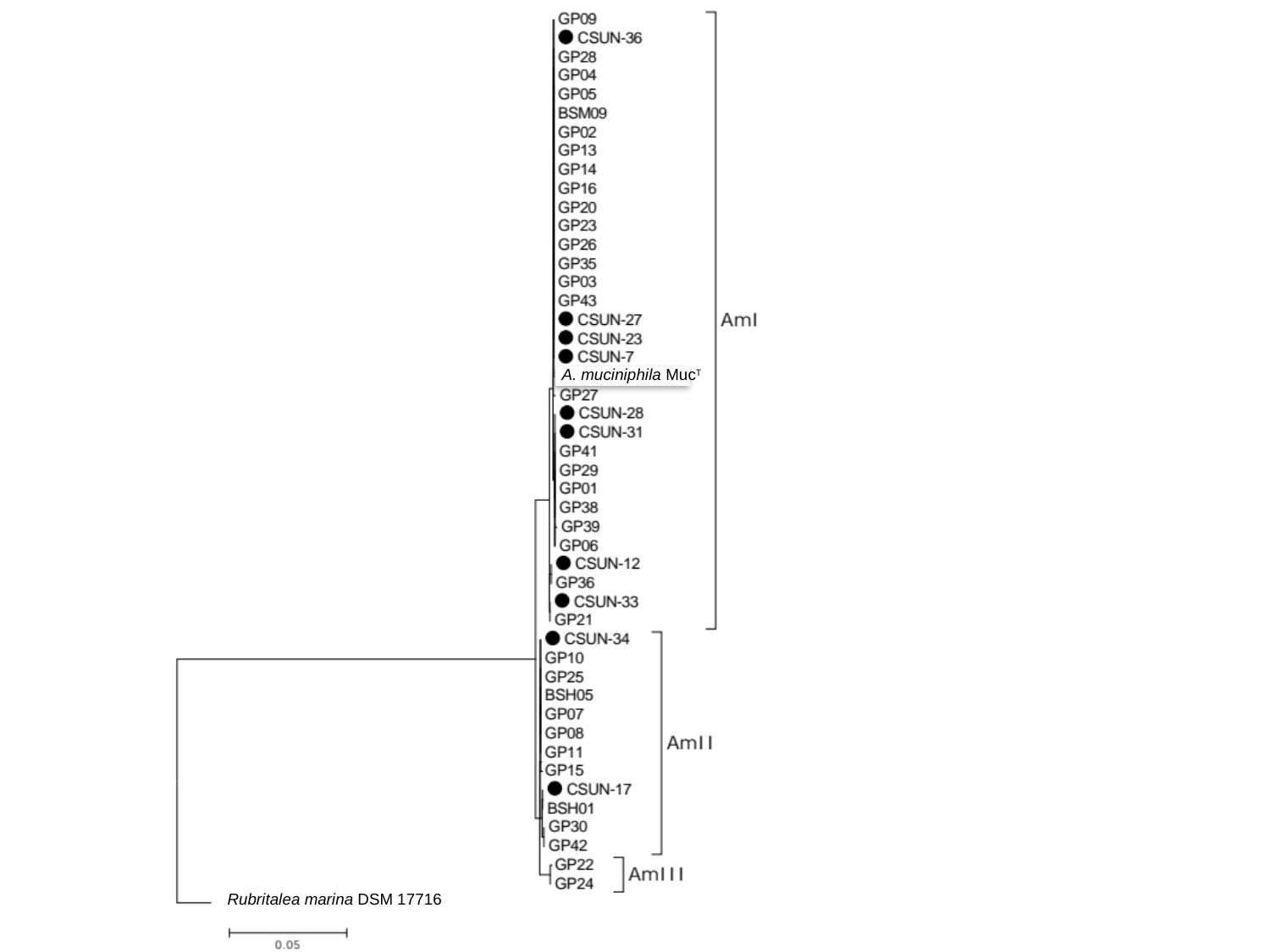

Rubritalea marina DSM 17716
A. muciniphila Muc^T^
